# Reproducible Data Analysis Pipelines for Precision Medicine

**DOI:** 10.1101/354811

**Authors:** Bjørn Fjukstad, Vanessa Dumeaux, Michael Hallett, Lars Ailo Bongo

## Abstract

Precision medicine brings the promise of more precise diagnosis and individualized therapeutic strategies from analyzing a cancer’s genomic signature. Technologies such as high-throughput sequencing enable cheaper data collection at higher speed, but rely on modern data analysis platforms to extract knowledge from these high dimensional datasets. Since this is a rapidly advancing field, new diagnoses and therapies often require tailoring of the analysis. These pipelines are therefore developed iteratively, continuously modifying analysis parameters before arriving at the final results. To enable reproducible results it is important to record all these modifications and decisions made during the analysis process.

We built a system, `walrus`, to support reproducible analyses for iteratively developed analysis pipelines. The approach is based on our experiences developing and using deep analysis pipelines to provide insights and recommendations for treatment in an actual breast cancer case. We designed `walrus` for the single servers or small compute clusters typically available for novel treatments in the clinical setting. `walrus` leverages software containers to provide reproducible execution environments, and integrates with modern version control systems to capture provenance of data and pipeline parameters.

We have used `walrus` to analyze a patient’s primary tumor and adjacent normal tissue, including subsequent metastatic lesions. Although we have used `walrus` for specialized analyses of whole-exome sequencing datasets, it is a general data analysis tool that can be applied in a variety of scientific disciplines. We have open sourced `walrus` along with example data analysis pipelines at github.com/uit-bdps/walrus.

## 1 Introduction

Precision medicine uses patient-specific molecular information to diagnose and categorize disease to tailor treatment to improve health outcome.[1] Important goals in precision medicine are to learn about the variability of the molecular characteristics of individual tumors, their relationship to outcome, and to improve diagnosis and therapy.[2] International cancer institutions are therefore offering dedicated personalized medicine programs.

For cancer, high throughput sequencing is an emerging technology to facilitate personalized diagnosis and treatment since it enables collecting high quality genomic data from patients at a low cost. Data collection is becoming cheaper, but the downstream computational analysis is still time consuming and thereby a costly part of the experiment. This is because of the manual efforts to set up, analyze, and maintain the analysis pipelines. These pipelines consist of a large number of steps that transform raw data into interpretable results.[3] These pipelines often consists of in-house or third party tools and scripts that each transform input files and produce some output. Although different tools exist, it is necessary to carefully explore different tools and parameters to choose the most efficient to apply for a dedicated question.[4] The complexity of the tools vary from toolkits such as the Genome Analysis Toolkit (GATK) to small custom *bash* or *R* scripts. In addition some tools interface with databases whose versions and content will impact the overall result.[5]

Improperly developed analysis pipelines for precision medicine may generate inaccurate results, which may have negative consequences for patient care.[6] When developing analysis pipelines for use in precision medicine it is therefore necessary to track pipeline tool versions, their input parameters, and data. Both to thoroughly document what produced the final clinical reports, but also for iteratively improving the quality of the pipeline during development. Because of the iterative process of developing the analysis pipeline, it is necessary to use analysis tools that facilitate modifying pipeline steps and adding new ones with little developer effort.

### 1.1 Breast Cancer Diagnosis and Treatment

We have previously analyzed DNA sequence data from a breast cancer patient’s primary tumor and adjacent normal cells to identify the molecular signature of the patient’s tumor and germline. When the patient later relapsed we analyzed sequence data from the patient’s metastasis to provide an extensive comparison against the primary and to identify the molecular drivers of the patient’s tumor.

We used Whole-Genome Sequencing (WGS) to sequence the primary tumor and adjacent normal cells at an average depth of 20, and Whole-Exome Sequencing (WES) at an average depth of 300. The biological samples were sequenced at the Genome Quebec Innovation Centre and we stored the raw datasets on our in-house server. From the analysis pipelines we generated reports with end results, such as detected somatic mutations, that was distributed to both the patient and the treating oncologists. These could be used to guide diagnosis and treatment, and give more detailed insight into both the primary and metastasis. When the patient relapsed we analyzed WES data using our own pipeline manager, `walrus`, to investigate the metastasis and compare it to the primary tumor.

For the initial WGS analysis we developed a pipeline to investigate somatic and germline mutations based on Broad Institute’s best practices. We developed the analysis pipeline on our in-house compute server using a *bash* script version controlled with *git* to track changes as we developed the analysis pipeline. The pipeline consisted of tools including picard,^1^ fastqc,^2^ trimmomatic,^3^ and the GATK.^4^ While the analysis tools themselves provide the necessary functionality to give insights in the disease, ensuring that the analyses could be fully reproduced later left areas in need of improvement.

We chose a command-line script over more complex pipelining tools or workbenches such as Galaxy[7] because of its fast setup time on our available compute infrastructure, and familiar interface. More complex systems could be beneficial in larger research groups with more resources to compute infrastructure maintenance, whereas command-line scripting languages require little infrastructure maintenance over normal use. In addition, while there are off-site solutions for executing scientific workflows, analyzing sensitive data often put hard restrictions on where the data can be stored and analyzed.

After we completed the first round of analyses we summarized our efforts and noted some lessons learned.

- Datasets and databases should be version controlled and stored along with the pipeline description. In the analysis script we referenced to datasets and databases by their physical location on a storage system, but these were later moved without updating the pipeline description causing extra work. A solution would be to add the data to the same version control repository hosting the pipeline description.
- The specific pipeline tools should also be kept available for later use. Since installing many bioinformatics tools require a long list of dependencies, it is beneficial to store the pipeline tools to reduce the time to start analyzing new data or re-run analyses.
- It should be easy to add new tools to an existing pipeline and execution environment. This includes installing the specific tool and adding to an existing pipeline. Bundling tools within software containers, such as Docker, and hosting them on an online registry simplifies the tool installation process since the only requirement is the container runtime.
- While bash scripts have their limitations, using a well-known format that closely resembles the normal command-line use clearly have its advantages. It is easy to understand what tools were used, their input parameters, and the data flow. However, from our experience when these analysis scripts grow too large they become too complex to modify and maintain.
- While there are new and promising state-of-the art pipeline managers, many of these also require state-of-the-art computing infrastructure to run. This may not be the case for the current research and hospital environments.

The above problem areas are not just applicable to our research group, but common to other research and precision medicine projects as well. Especially when hospitals and research groups aim to apply personalized medicine efforts to guide therapeutic strategies and diagnosis, the analyses will have to be able to be easily reproducible later. We used the lessons learned to design and implement `walrus`, a command line tool for developing and running data analysis pipelines. It automatically orchestrates the execution of different tools, and tracks tool versions and parameters, as well as datasets through the analysis pipeline. It provides users a simple interface to inspect differences in pipeline runs, and retrieve previous analysis results and configurations. In the remainder of the paper we describe the design and implementation of `walrus`, its clinical use, its performance, and how it relates to other pipeline managers.

## 2 walrus

`walrus` is a tool for developing and executing data analysis pipelines. It stores information about tool versions, tool parameters, input data, intermediate data, output data, as well as execution environments to simplify the process of reproducing data analyses. Users write descriptions of their analysis pipelines using a familiar syntax and `walrus` uses this description to orchestrate the execution of the pipeline. In `walrus` we package all tools in software containers to capture the details of the different execution environments. While we have used `walrus` to analyse high-throughput datasets in precision medicine, it is a general tool that can analyze any type of data, e.g. image datasets for machine learning. It has few dependencies and runs on on any platform that supports Docker containers. While other popular pipeline managers require the use of cluster computers or cloud environment, we focus on single compute nodes often found in clinical environments such as hospitals.

`walrus` is implemented as a command-line tool in the Go programming language. We use the popular software container implementation Docker^5^ to provide reproducible execution environments, and interface with git together with git-lfs^6^ to version control datasets and pipeline descriptions. By choosing Docker and git we have built a tool that easily integrates with current bioinformatic tools and workflows. It runs both natively or within its own Docker container to simplify its installation process.

With `walrus` we target pipeline developers that use command-line tools and scripting languages to build and run analysis pipelines. Users can use existing Docker containers from sources such as BioContainers,[8] or build containers with their own tools. We integrate with the current workflow using git to version control analysis scripts, and use git-lfs for versioning of datasets as well. We have designed the pipeline description format resembles the command line syntax as much as possible. This is one of the major strengths of walrus. It uses a familiar syntax and format, and does not require the users to explicitly declare which files in the pipeline to version control.

### 2.1 Pipeline Configuration

Users configure analysis pipelines by writing pipeline description files in a human readable format such as JavaScript Object Notation (JSON) or YAML Ain’t Markup Language (YAML). A pipeline description contains a list of stages, each with inputs and outputs, along with optional information such as comments or configuration parameters such as caching rules for intermediate results. Listing 1 shows an example pipeline stage that uses MuTect[9] to detect somatic point mutations. Users can also specify the tool versions by selecting a specific Docker image, for example using MuTect version 1.1.7 as in Listing 1, line 3.

Users specify the flow of data in the pipeline within the pipeline description, as well as the dependencies between the steps. Since pipeline configurations can become complex, users can view their pipelines using an interactive web-based tool, or export their pipeline as a DOT file for visualization in tools such as Graphviz.^7^

Listing 1: Example pipeline stage for a tool that detects somatic point mutations. It reads a reference sequence file together with both tumor and normal sequences, and produces an output file with the detected mutations.

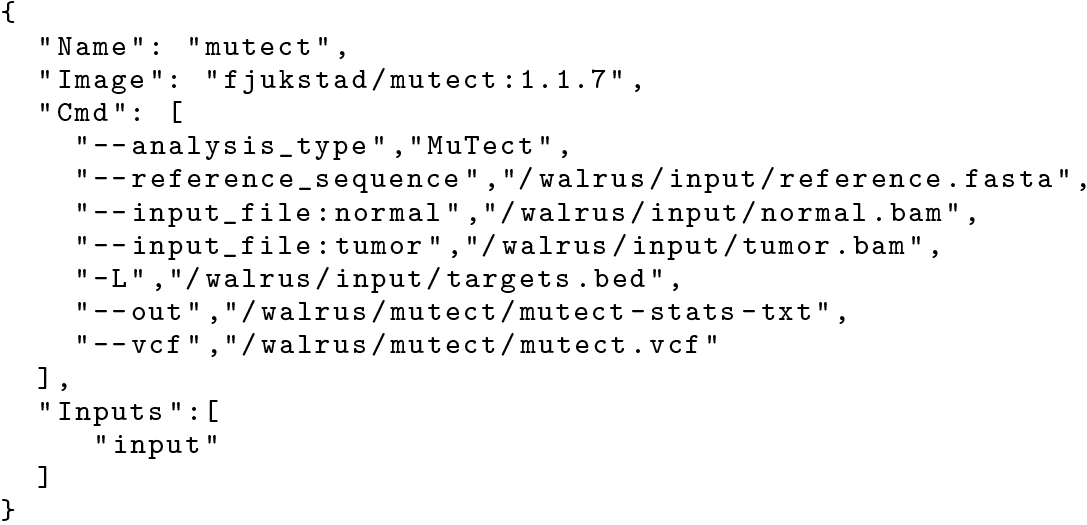

Users add data to an analysis pipeline by specifying the location of the input data in the pipeline description, and `walrus` automatically mounts it to the container running the analysis. The location of the input files can either be local or remote locations such as an FTP server. When the pipeline is completed, `walrus` will store all the input, intermediate and output data to a user-specified location.

### 2.2 Pipeline Execution

When users have written a pipeline description for their analyses, they can use the command-line interface of `walrus` to run the analysis pipeline. `walrus` builds an execution plan from the pipeline description and runs it for the user. It uses the input and output fields of each pipeline stage to construct a Directed Acyclic Graph (DAG) where each node is a pipeline stage and the links are input/output data to the stages. From this graph `walrus` can determine parallelizable stages and coordinate the execution of the pipeline.

In `walrus`, each pipeline stage is run in a separate container, and users can specify container versions in the pipeline description to specify the correct version of a tool. We treat a container as a single executable and users specify tool input arguments in the pipeline description file using standard command line syntax. `walrus` will automatically build or download the container images with the analysis tools, and start these with the user-defined input parameters and mount the appropriate input datasets. While the pipeline is running, `walrus` monitors running stages and schedules the execution of subsequent pipeline stages when their respective input data become available. We have designed `walrus` to execute an analysis pipeline on a single large server, but since the tools are run within containers, these can easily be orchestrated across a range of servers in future versions.

Users can select from containers pre-installed with bioinformatics tools, or build their own using a standard Dockerfile. Through software containers `walrus` can provide a reproducible execution environment for the pipeline, and containers provide simple execution on a wide range of software and hardware platforms. With initiatives such as BioContainers, researchers can make use of already existing containers without having to re-write their own. Data in each pipeline step is automatically mounted and made available within each Docker container. By simply relying on Docker `walrus` requires little software setup to run different bioinformatics tools.

While `walrus` executes a single pipeline on one physical server, it supports both data and tool parallelism, as well as any parallelization strategies within each tool, e.g. multi-threading. If users want to run the same analyses for a set of samples, or for example per chromosome, they can simply list the samples in the pipeline description and `walrus` will automatically run each sample through the pipeline in parallel. While we can parallelize the independent pipeline steps, the performance of an analysis pipeline relies on each of the independent tools and available compute power. This also applies to the scalability of the analysis pipeline.

Upon successful completion of a pipeline run, `walrus` will write a verbose pipeline description file to the output directory. This file contains information on the runtime of each step, which steps were parallelized, and provenance related information to the output data from each step. Users can investigate this file to get a more detailed look on the completed pipeline. In addition to this output file `walrus` will return a unique version ID for the pipeline run, which later can be used to investigate a previous pipeline run.

### 2.3 Data Management

In `walrus` we provide an interface for users to track their analysis data through a version control system. This allows users to inspect data from previous pipeline runs without having to recompute all the data. `walrus` stores all intermediate and output data in an output directory specified by the user, which is version controlled automatically by `walrus` when new data is produced by the pipeline. We track changes at file granularity.

In `walrus` we interface with `git` to track any output file from the analysis pipeline. When users execute a pipeline, `walrus` will automatically add and commit output data t a git repository using `git-lfs`. Users typically use a single repository per pipeline, but can share the same repository to version multiple pipelines as well. With git-lfs, instead of writing large blobs to a repository it writes small pointer files that contains the hash of the original file, the size of the file, and the version of git-lfs used. The files themselves are stored separately which makes the size of the repository small and manageable with git. The main reason why we chose git and git-lfs for version control is that git is the de facto standard for versioning source code, and we want to include versioning of datasets without altering the typical development workflow.

Since we are working with potentially sensitive datasets `walrus` is targeted at users that use a local compute and storage servers. It is up to users to configure a remote tracker for their repositories, but we provide command-line functionality in `walrus` to run a git-lfs server that can store users’ contents. They can use their default remotes, such as Github, for hosting source code but they must themselves provide the remote server to host their data.

### 2.4 Pipeline Reconfiguration and Re-execution

Reconfiguring a pipeline is common practice in precision medicine, e.g. to ensure that genomic variants are called with a desired sensitivity and specificity. To reconfigure an existing pipeline users make the applicable changes to the pipeline description and re-run it using `walrus`. `walrus` will then recompute the necessary steps and return a version ID for the newly run pipeline. This ID can be used to compare pipeline runs, the changes made, and optionally restore the data and configuration from a previous run. Reconfiguring the pipeline to use updated tools or reference genomes will alter the pipeline configuration and force `walrus` to recompute the applicable pipeline stages.

**Figure 1:**
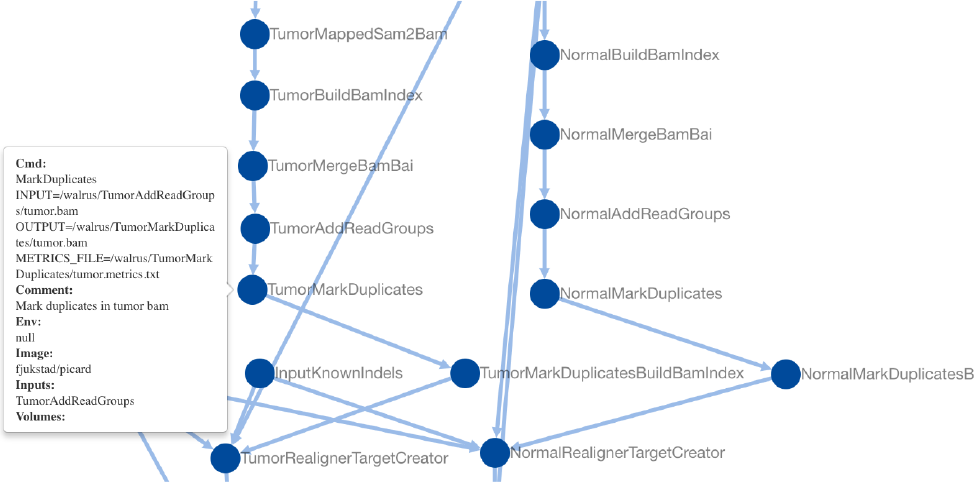
Screenshot of the web-Based visualization in `wairus`. The user has zoomed in to inspect the pipeline step which marks duplicate reads in the tumor sequence data.

The command-line interface of `walrus` provides functionality to restore results from a previous run, as well as printing information about a completed pipeline. To restore a previous pipeline run, users use the `restore` command line flag in `walrus` together with the version ID of the respective pipeline run. `walrus` will interface with git to restore the files to their state at the necessary point in time.

## 3 Results

To evaluate the usefulness of `walrus` we demonstrate its use in a clinical setting, and the low computational time and storage overhead to support reproducible analyses.

### 3.1 Clinical Application

We have used `walrus` to analyze a whole-exome data from a sample in the McGill Genome Quebec [MGGQ] dataset (GSE58644)[10] to discover Single Nucleotide Polymorphisms (SNPs), genomic variants and somatic mutations. We interactively developed a pipeline description that follows the best-practices of The Broad Institute^7^ and generated reports that summarized the findings to share the results. Figure 1 shows a screenshot from the web-based visualization in `walrus` of the pipeline.

From the analyses we discovered inherited germline mutations that are recognized to be among the top 50 mutations associated with an increased risk of familial breast cancer. We also discovered a germline deletion which has been associated with an increased risk of breast cancer. We also discovered mutations in a specific gene that might explain why specific drug had not been effective in the treatment of the primary tumor. From the profile of the primary tumor we discovered many somatic events (around 30 000) across the whole genome with about 1000 in coding regions, and 500 of these were coding for non-synonymous mutations. We did not see amplification or constituent activation of growth factors like HER2, EGFR or other players in breast cancer. Because of the germline mutation, early recurrence, and lack of DNA events, we suspect that the patient’s primary tumor was highly immunogenic. We have also identified several mutations and copy number changes in key driver genes. This includes a mutation in a gene that creates a premature stop codon, truncating one copy of the gene.

While we cannot share the results in details or the sensitive dataset, we have made the pipeline description available at github.com/uit-bdps/walrus along with other example pipelines.

### 3.2 Example Dataset

To demonstrate the performance of `walrus` and the ability to track and detect changes in an analysis pipeline, we have implemented one of the variant calling pipelines from [11] using tools from picard and the GATK. We show the storage and computational overhead of our approach, and the benefit of capturing the pipeline specification using a pipeline manager rather than a methods section in a paper. The pipeline description and code is available along with `walrus` at github.com/uit-bdps/walrus. Figure 2 shows a simple graphical representation of the pipeline.

#### 3.2.1 Performance and Resource Usage

We first run the variant calling pipeline without any additional provenance tracking or storing of output or intermediate datasets. This is to get a baseline performance measurement for how long we expect the pipeline to run. We then run a second experiment to measure the overhead of versioning output and intermediate data. Then we introduce a parameter change in one of the pipeline steps which results in new intermediate and output datasets. Specifically we change the `--maxReadsForRealignment` parameter in the indel realigner step back to its default (See the online pipeline description for more details). This forces `walrus` to recompute the indel realigner step and any subsequent steps. We then use the `restore` flag in `walrus` to illustrate what the parameter change had on the pipeline output. To illustrate how `walrus` can restore old pipeline configurations and results, we restore the pipeline to the initial configuration and results. We show the computational overhead and storage usage of restoring a previous pipeline configuration.

Reproducing results from a scientific publication can be a difficult task. For example, troublesome formatting of the online version of [11] led to some pipeline tools failing. The parameters prefixed with two consecutive hypens (--) are converted to single em dashes (—). PDF versions of the paper lists the parameters correctly. In addition, the input filenames in the variant calling step do not correspond to any output files in previous steps, but because of their similarity to previous output files we assume that this is just a typo. These issues in addition to missing commands for e.g. the filtering step highlights the clear benefit of writing and reporting the analysis pipeline using a tool such as `walrus`.

**Figure 2:**
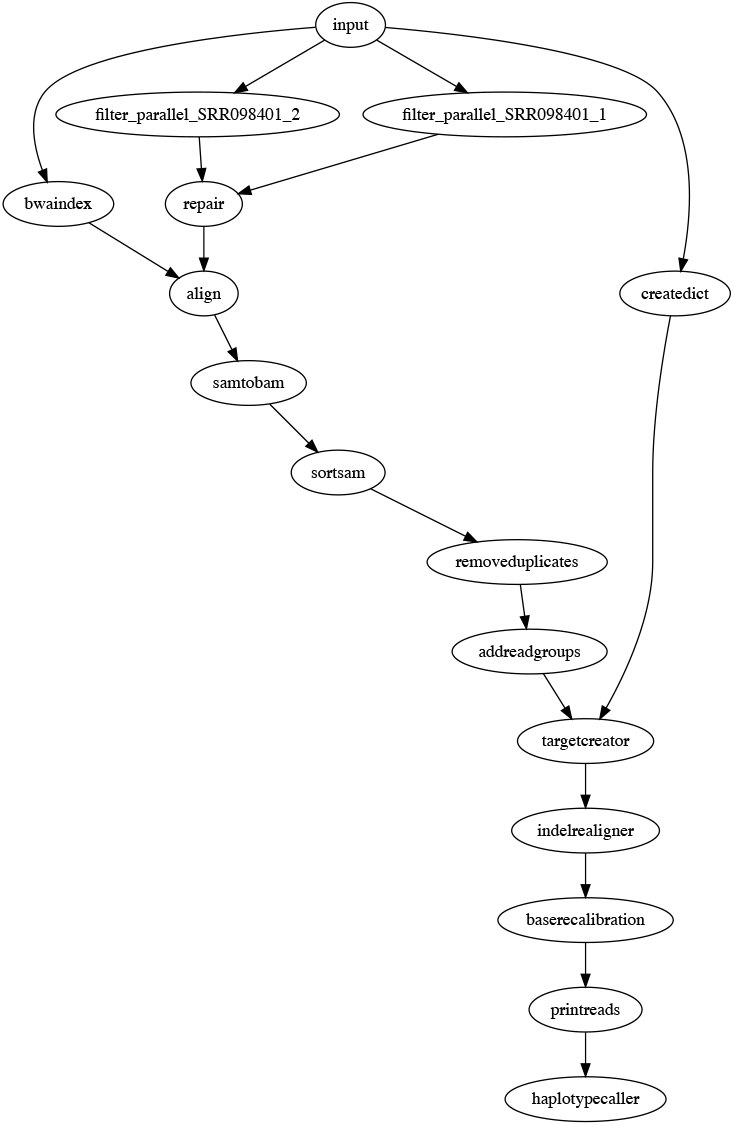
In addition to the web-based inteactive pipeline visualization, `walrus` can also generate DOT representations of pipelines. The figure shows the example variant calling pipeline.

Table 1 shows the runtime and storage use of the different experiments. In the second experiment we can see the added overhead of adding version control to the dataset. In total, an hour is added to the runtime and the data size is doubled. The doubling comes from git-lfs hard copying the data into a subdirectory of the `.git` folder in the repository. With git-lfs users can move all datasets to a remote server reducing the local storage requirements. In the third experiment we can see that only the downstream analyses from configuring the indel realignment parameter is executed. It generates 30GB of additional data, but the execution time is limited to the applicable stages. Restoring the pipeline to a previous configuration is almost instantaneous since the data is already available locally and git only has to modify the pointers to the correct files in the `.git` subdirectory.

**Table 1:**
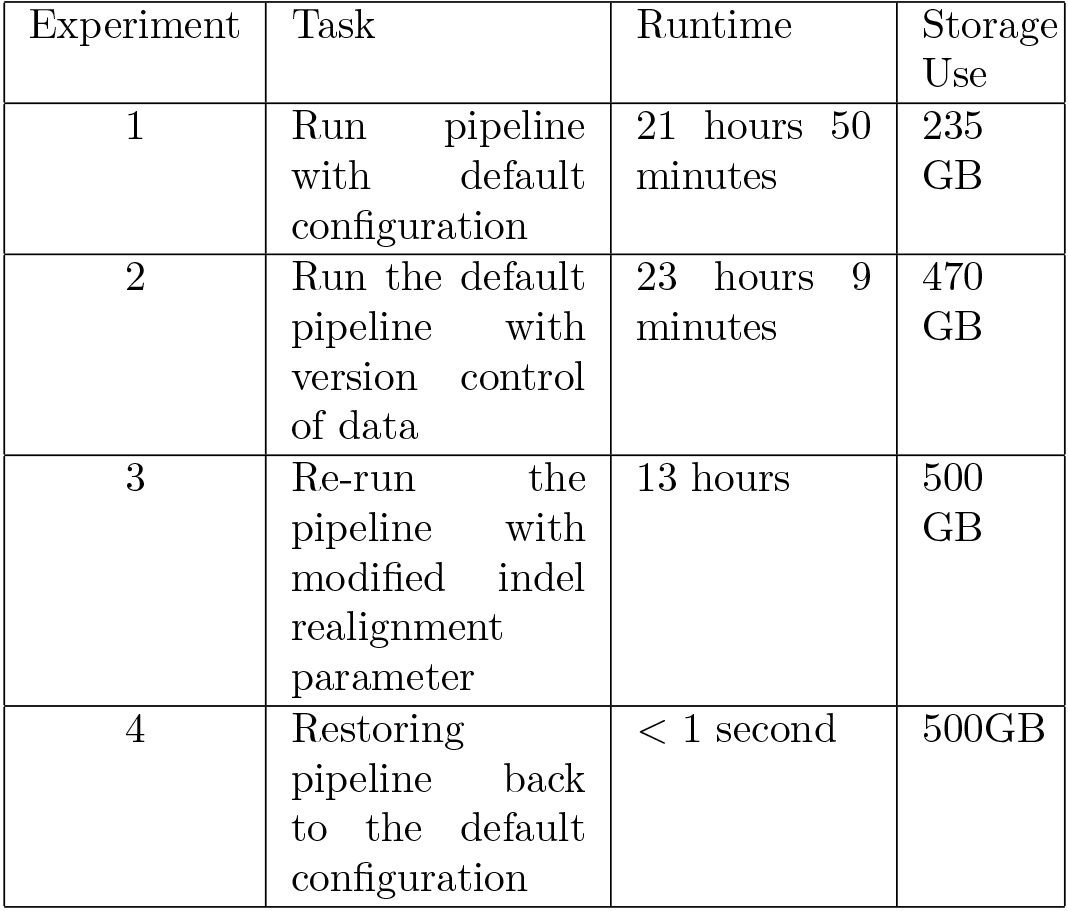
Runtime and storage use of a the typical workflow of developing a variant-calling pipeline with `walrus`.

## 4 Related Work

There are a wealth of pipeline specification formats and workflow managers available. Some are targeted at users with programming experience while others provide simple Graphical User Interfaces (GUIs). Here we describe the most popular systems for building data analysis pipelines. While most provide viable options for genomic analyses, we have found most to complex to install and maintain in clinical settings. We discuss tools that use the common Common Workflow Language (CWL) pipeline specification and systems that provide versioning of data.

CWL is a specification for describing analysis workflows and tools.[12] A pipeline is written as a JSON or YAML file, or a mix of the two, and describes each step in detail, e.g. what tool to run, its input parameters, input data and output data. The pipeline descriptions are text files that can be version controlled and shared between projects. There are multiple implementations of CWL workflow platforms, e.g. the reference implementation cwl_runner,^9^ Arvados,[13] Rabix,[14] Toil,[15] Galaxy,[7] and AWE.[16] It is no requirement to run tools within containers, but implementations can support it. There are few of these tools that support versioning of the data. Galaxy is an open web-based platform for reproducible analysis of large high-throughput datasets.[7] It is possible to run Galaxy on local compute clusters, in the cloud, or using the online Galaxy site.^9^ In Galaxy users set up an analysis pipeline using a web-based graphical interface, and it is also possible to export or import an existing workflow to an Extensible Markup Language (XML) file.^11^ We chose not to use Galaxy because of missing command-line and scripting support, along with little support for running workflows with different configurations.[17] Rabix provides checksums of output data to verify it against the actual output from the pipeline. This is similar to the checksums found in the git-lfs pointer files, but they do not store the original files for later. Arvados stores the data in a distributed storage system, Keep, that provides both storage and versioning of data. We chose not to use CWL and its implementations because of its relaxed restrictions on having to use containers, its verbose pipeline descriptions, and the complex compute architecture required for some of the runners. We are however experimenting with an extension to `walrus` that translates pipeline descriptions written in `walrus` to CWL pipeline descriptions.

Pachyderm is a system for running big data analysis pipelines. It provides complete version control for data and leverages the container ecosystem to provide reproducible data processing.^12^ Pachyderm consists of a file system (Pachyderm File System (PFS)) and a processing system (Pachyderm Processing System (PPS)). PFS is a file system with git-like semantics for storing data used in data analysis pipelines. Pachyderm ensures complete analysis reproducibility by providing version control for datasets in addition to the containerized execution environments. Both PFS and PPS is implemented on top of Kubernetes.^13^ We believe that the approach in Pachyderm with version controlling datasets and containerizing each pipeline step is the correct approach to truly reproducible data analysis pipelines. The reason we did not use Kubernetes and Pachyderm was because our compute infrastructure did not support it. In addition we did not want to use a separate tool, PFS, for data versioning, we wanted to integrate it with the current pratice of using git for versioning.

The BioContainers[8] and Bioboxes[18] projects address the challenge of installing bioinformatics data analysis tools by maintaining a repository of Docker containers for commonly used data analysis tools. Docker containers are shown to have better than, or equal performance as VMs.[19] Both forms of virtualization techniques introduce overhead in I/O-intensive workloads, especially in VMs, but introduce negligible CPU and memory overhead. For precision medicine pipelines the overhead of Docker containers will be negligible since these tend to be compute intensive and they typically run for several hours. [19] Containers have also been proposed as a solution to improve experiment reproducibility, by ensuring that the data analysis tools are installed with the same responsibilities.[20]

## 5 Discussion

Precision medicine requires flexible analysis pipelines that allow researchers to explore different tools and parameters to analyze their data. While there are best practices to develop analysis pipelines for genomic datasets, e.g. to discover genomic variants, there is still no de-facto standard for sharing the detailed descriptions to simplify re-using and reproducing existing work. Pipelines typically need to be tailored to fit each project and patient, and different patients will typically elicit different molecular patterns that require individual investigation. While we could follow best practices to develop our pipeline we explored different tools and parameters before we arrived at the final analysis pipeline. For example, in our WES pipeline we ran several rounds of preprocessing (trimming reads and quality control) before we were sure that the data was ready for analysis. Having a pipeline system that could keep track of different intermediate datasets, along with the pipeline specification, simplifies the task of comparing the results from pipeline tools and input parameters. While we have developed one approach to version control genomic datasets in an analysis pipeline, we believe that there is still room for improvement.

While we provide one approach to version control datasets, there are still some drawbacks. git-lfs supports large files, but in our results it added an additional 5% in runtime. This makes the entire analysis pipeline slower, but we argue that having the files version controlled outweigh the runtime. In addition, there are only a few public gif-lfs hosting platforms for datasets larger than a few gigabytes, making it necessary to host these in house.

We aim to investigate the performance of running analysis pipelines with `walrus`, and the potential benefit of its built-in data parallelism. While our WES analysis pipeline successfully run steps in parallel for the tumor and adjacent normal tissue, we have not demonstrated the benefit of doing so. This includes benchmarking and analyzing the system requirements for doing precision medicine analyses. We are also planning on exploring parallelism strategies where we can split an input dataset into chromosomes and run some steps in parallel for each chromosome, before merging the data again.

## 6 Conclusions

We have designed and implemented `walrus`, a tool for developing reproducible data analysis pipelines for use in precision medicine. Precision medicine requires that analyses are run on hospital compute infrastructures and results are fully reproducible. By packaging analysis tools in software containers, and tracking both intermediate and output data, `walrus` provides the foundation for reproducible data analyses in the clinical setting. We have used `walrus` to analyze a patient’s metastatic lesions and adjacent normal tissue to provide insights and recommendations for cancer treatment.

## 7 Acknowledgements

We would like to thank Daniel Del Balso for his work implementing the initial WGS analysis pipeline.

This work has been funded by The European Research Council (ERC-AdG 232997 TICE), and The Canadian Cancer Society Research Institute (INNOV2-2014-702940).

## Footnotes

1 broadinstitute.github.io/picard

2 bioinformatics.babraham.ac.uk/projects/fastqc

3 usadellab.org/cms/?page=trimmomatic

4 software.broadinstitute.com/gatk

5 docker.com

6 git-lfs.github.com

7 graphviz.org

8 software.broadinstitute.org/gatk/best-practices

9 github.com/common-workflow-language/cwltool

10 Availableatusegalaxy.org.

11 An alpha version of Galaxy with CWL support is available at github.com/common-workflow-language/galaxy.

12 pachyderm.io

13 kubernetes.io

